# Plant allometry derived from Metabolic Scaling Theory and segregated by tissue functionality

**DOI:** 10.1101/2023.08.10.552736

**Authors:** S. Sopp, R. Valbuena

## Abstract

Plant allometry is key for determining the role of forests in global carbon cycles, through the calculation of tree biomass using proxy measurements such as tree diameters or heights. Metabolic Scaling Theory (MST) considers the general principles that underpin allometry, but MST scaling relationships have been challenged on their lack of fit to empirical data and global applicability. Many authors have thus optimised their model forms for statistical performance over theory based approaches. We postulated that MST scaling is applicable only to the proportion of plant tissue with supportive functionality, meaning that as plants evolved tissues of specialized conductive functionality (e.g vessels) their allometry progressed into more complex relationships. Our generalised MST (gMST) models were thus created by considering conductive lumen as unsupportive area, and consequentially removing it from the original MST 2/3 scaling. According to this principle, we deducted generalized gMST relationships with mechanistically deducted coefficients. We found that the gMST height-diameter scaling outperformed current state of the art equations that are widely used within the tropics and that the model performed well across all tested ecoregions. Furthermore, the proposed aboveground biomass models performed similarly to widely used models in the literature within the tropics. The results presented indicate that the further development of generalised allometric models remains a research priority given the importance of assessing and monitoring global forest carbon fluxes. The height-diameter models presented can thus be of much use to the wider community in further refining carbon stock estimates globally, providing a universally applicable theoretical framework.

## Introduction

The assessment and monitoring of carbon fluxes within the biosphere is paramount now more so than ever, due to the capacity of forests to help mitigate climate change through programs like REDD+ (Corbera & Schroeder, 2011; Gibbs et al., 2007). The ability to accurately and reliably estimate forest carbon stocks is vital to this endeavour, meaning that well adjusted plant scaling models are required. The need for statistically powerful allometric equations at large scales therefore continues to be an important area of research (Jucker et al., 2022, 2017; Chave et al., 2014). To this end a plethora of allometric equations exist within the literature, with a wide variety of model forms hosted across a vast geographic and taxonomic scope (Chojnacky et al., 2014; Zianis et al., 2005; Chave et al., 2014; Zhou et al., 2021; Valbuena et al., 2016). Many of these equations are however not based upon allometric theory, and instead select model form based upon statistical performance, like that of Chave et al. (2014), whose models are widely used within the tropics. Such optimised models unsurprisingly tend to outperform theory based models like that of Metabolic Scaling Theory (MST) (West et al., 1999). Consequentially theory based allometric models are seldom used for large scale carbon stock estimates.

Metabolic scaling theory (MST) represents the pinnacle of theory-based understanding for biological scaling, providing a broad framework of allometric equations (West et al., 1997, 1999). The theory has implications for understanding ecosystem growth and dynamics (Enquist et al., 2000), ecosystem structure (Enquist et al., 2009; K. Niklas & Enquist, 2001; Lee et al., 2021), and to a greater extent net primary productivity and ecosystem energy and carbon cycles (Allen et al., 2005; Price et al., 2010; Schramski et al., 2015). However, the plant allometric relationships derived through MST have faced criticism for their lack of empirical support, with larger trees particularly deviating from the theories predictions (Reich et al., 2006; Muller-Landau et al., 2006; D. A. Coomes, 2006). For this reason, there has been attempts to adjust its theoretical basis (Savage VM, 2008), such as the inclusion of additional aspects like canopy geometry packing dynamics (Taubert et al., 2015; Lutz et al., 2018) or the influence of asymmetric competition of larger trees for resources (D. Coomes et al., 2011). Despite these attempts, and the general revision of the theory’s vascular underpinning (Sperry et al., 2006; Savage et al., 2010), a mechanistic advancement that explains the lack of fit for large diameter trees not been realised.

Some of the key allometric equations that stem from MST provide scaling relationships between diameter (*d*) and height (*h*), volume (*v*_*tot*_) or aboveground biomass (*agb*) and subsequently carbon. MST (West et al., 1999) dictates that the *h* an individual tree reaches scales to 2/3 of its *d*. However this *h* -*d* relationship often fits poorly to real tree data, with many studies fitting power laws notably below 2/3 (Muller-Landau et al., 2006; Picard et al., 2015; Swetnam et al., 2016; Jucker et al., 2017), with the greatest deviation shown by large trees which exhibit exponents significantly lower (K. J. Niklas, 1995; Muller-Landau et al., 2006; Banin et al., 2012). Consequently, the geometrically propagated *agb* -*d* exponent of 8/3 predicted by MST (West et al., 1999) is consistently higher than adjusted models worldwide (Aabeyir et al., 2020; Zianis & Mencuccini, 2004; Zianis et al., 2005; Chojnacky et al., 2014). The exponents predicted by MST are well established within the literature, however the consistent and significant deviation from MST exponents signifies a lack of mechanistic understanding. Scholars have therefore taken to using model forms other than power laws that empirically suit their datasets (Chave et al., 2014; Barbosa et al., 2019; Valbuena et al., 2016). Statistically optimised model forms have therefore been used preferentially over theory-based models (Picard et al., 2015; Feldpausch et al., 2012; Zhou et al., 2021). The question is therefore raised as to whether a suitable mechanistic model other than a simple power relationship can be derived from MST, conciliating the current divide between theoretical and statistical plant allometry models.

Hydraulic and mechanical constraints are typically cited as key influences of plant allometry, with their interaction governing *h* (K. Niklas, 2007). A trade-off between these two constraints is evident (Sperry et al., 2008), particularly for large trees (Long et al., 1981; Zhou et al., 2021). Each constraint can be considered through the proportion of area with conductive (*CA*) and supportive (*SA*) functionality within a cross-section of the stem (see Fig. 1 below)(Ziemińska et al., 2013; Zanne et al., 2010). At a cellular level, these functions manifest differently between species, as angiosperms often contain vessels with dedicated lumen for fluid conductance and fibres for support. Alternatively gymnosperms typically have tracheids that integrate both functions within one cell type, with conduits located within tracheids for fluid transport (Sperry et al., 2006). *CA* and *SA* are thus best considered as the proportional summation of conductive lumen area, and the remaining proportion of non-conductive lumen area that acts to support the plant respectively.

**Figure 1:**
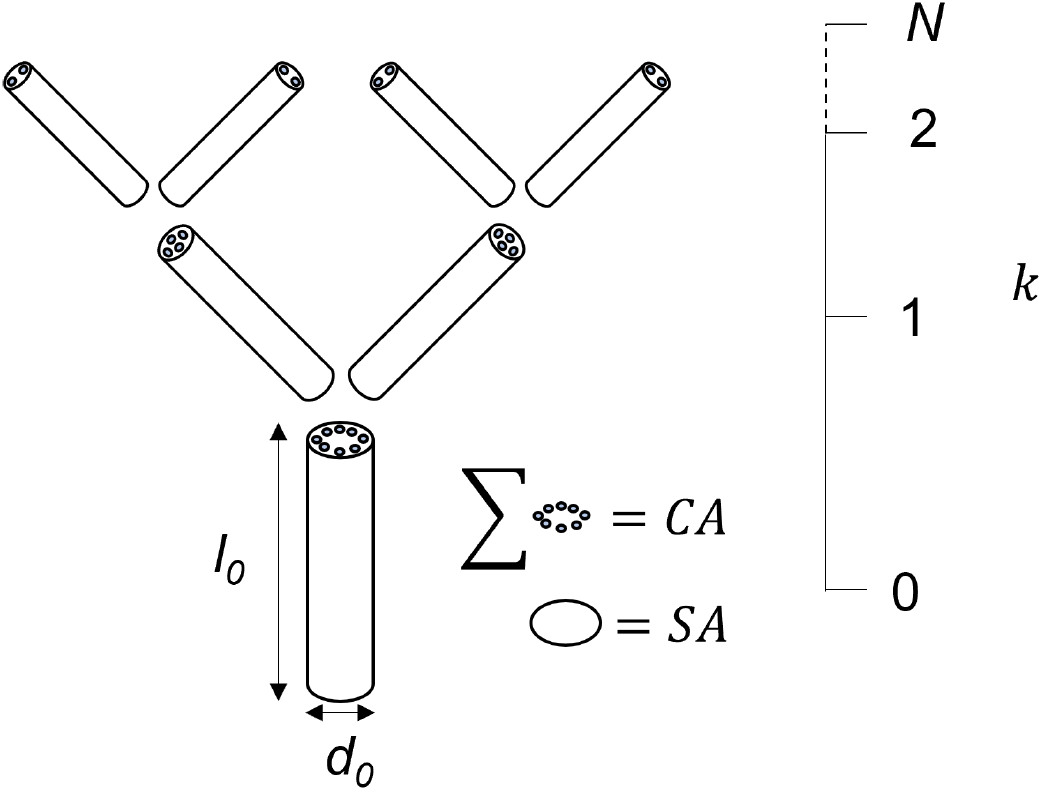
Illustration of a simple branching network with 3 levels given (0,1, and 2), as well as a depiction of vessels within the network. The summation of vessels/conduits divided by total area gives *CA*, while *SA* can be found through 1 − *CA. l*_0_ gives the length of the first generation in the branching network (aka the trunk) and *d*_0_ gives the diameter of the first branching generation (thus diameter of the trunk).

We postulated that a distinction between *CA* and *SA* is required to advance the current theory and overcome the shortfalls of the original MST scaling. We therefore revisited MST’s underpinning assumptions and used them to derive a new set of allometric equations, considering the interaction between conductive and supportive functionality. The mechanical and hydraulic limitations can thus be integrated into MST. We hypothesized that this advancement would generate a set of generalized MST (gMST) model forms for the *h* -*d* and *agb* -*d* relationships which would adequately fit vascular plants independently of functional type (angiosperm/gymnosperm) or geographical location (ecoregion), clarifying the reasons for the apparent lack of empirical fit in large trees and for certain tree species. We show that the gMST equations demonstrate statistical and conceptual validity globally, and outperform or prove comparable to statistical models that are widely used as the gold standard for estimating biomass carbon in tropical forests (Chave et al., 2014). Most importantly, these results substantiate that the implications of MST in plant allometry hold true globally independently of plant size.

## Rationale for the generalized MST (gMST) model

The 2/3 scaling relationship between *d* and *h* predicted by MST derives strictly from the biomechanical limitations of an area-preserving and volume filling network of branching geometry (West et al., 1999; K. Niklas, 2007; Sperry et al., 2006; McMahon & Kronauer, 1976; Savage et al., 2010). The original formulation of allometry derived from MST assumes that the biomechanical capabilities of both *CA* and *SA* are equivalent to derive the 2/3 exponent in the *h* ∝ *d* ^2*/*3^ scaling (West et al., 1997, 1999). Vessel and conduit lumen however fail to reinforce plant structure, with significant negative correlations with mechanical variables (Ziemińska et al., 2015).

Conduit/vessel lumen (the component of *CA*), is functionally adapted for fluid transport (conductivity) rather than wood strength, which is evidenced by negative correlations between lumen fraction and Young’s modulus and wood density (Ziemińska et al., 2015) as well as positive correlations with conductivity (Ziemińska et al., 2013). Both Young’s modulus and wood density affect the max height of trees (Greenhill, 1881). These variables are interconnected yielding a scaling relationship, and thus fail to form a constant meaning stiffer stems are proportionally less strong (K. J. Niklas, 1993). The exclusion of these variables from allometric models are therefore likely to lower the accuracy of tree height predictions. Given that *CA* has a significant relationship with both variables it could be a valuable addition to allometric models (Ziemińska et al., 2013, 2015). Tree growth can be seen as a complex process, with the transition of conducting sapwood to heartwood causing varying degrees of local conductivity and density across the width of a stem (Longuetaud et al., 2017). The conductive area of a stem is thus located in the outer section of the trunk, resulting in trees having an outer conductive segment (see Fig. 1). Most plants therefore have a degree of hollowness towards their periphery making up *CA*, in contrast to the modelling of trees as solid cylinders (or the modelling of hollow plants (Kanahama & Sato, 2022)). We decided to test a model form that included *SA* as a direct modifier to plant *d*, with it also scaling to 2/3, yielding the equation *h* ∝ (*d* · *SA*)^2*/*3^. The scaling for *SA* was chosen as to create an effective diameter term within the equation, removing *CA* which negatively correlates with mechanical ability. This approach therefore contrasts solid cylinder approaches (Greenhill, 1881), and assumptions of uniform biomechanical limitations (West et al., 1999). *CA*’s inclusion within the equation further allows for differences between angiosperms and gymnosperms to be realised within the model, given that the former can typically reach larger conduit dimensions (Yang et al., 2022). The inclusion of *SA* in such a fashion could resolve discrepancies between the ability of MST to successful model *h* -*d* relationships across functional groups.

A novel gMST model can thus be derived for *h* purely as a function of *d*. 1− *CA* offers an alternative representation of *SA*, generating the equation *h* ∝ (*d* · (1 − *CA*))^2*/*3^, where *CA* can be predicted by MST. The scaling of *CA* has been subject to some debate since MST’s inception, due to the conceptual modelling of the vascular system. West et al. (1999) described the vascular system as a fixed number of tapered tubes creating a decreasing proportion of vessel lumen along the plants length, with the scaling *CA* ∝ *d* ^1*/*3^. A later revision of MST Savage et al. (2010) considered the tubes to furcate such that lumen area was maintained, *CA* ∝ *d* ^0^, while both assumed the network was energy efficient and area preserving. However, recent evidence suggests a transition between the two models, with West et al. (1999)’s original derivation being more applicable for much of the stem (Koçillari et al., 2021; Rosell & Olson, 2019; Lechthaler et al., 2019), while the revised scaling is likely more applicable towards the termination of the network. We have therefore opted to use West et al. (1999)’s original scaling for *CA* given it is likely suitable across the majority of the network, yielding a novel MST based height-diameter equation:

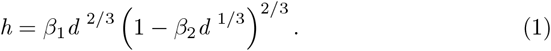

Like the original theory there are a series of assumptions that underpin this relationship, being area preservation and resistance maintenance. The assumption of area preservation is debated, with few studies considering the empirical strength of such an assumption. Those that have considered area preservation tend to find a small deviation from this assumption (Eloy, 2011), although not always statistically different from area preservation (Bentley et al., 2013). Thus we maintain the assumption of area preservation, if not only to simply the modelling procedure. Vascular optimality on the other hand appears to be broadly accepted within the literature (West et al., 1999; Savage et al., 2010; Koçillari et al., 2021) and is consequentially also assumed within our models.

With Eq. 1 established, a gMST based allometric *agb* relationship can be deduced. Based on MST assumptions tree *agb* can be given as a function of both its *d, h* and average wood density (*ρ*) (*agb* ∼ *f* (*d, h, ρ*)):

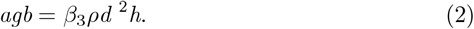

Eq. 2 implies that biomass is directly proportional to volume (i.e. an exponent equal to 1). Consequentially, the gMST *h* -*d* model form (Eq. 1) can be substituted into Eq. 2, giving a gMST *agb* -*d* model:

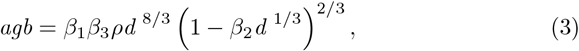

where *β*_1*−*3_ represent coefficients that can be mechanistically deducted through tree parameters established by MST and associated scalars (see Eqs. 8 and 11 in Methods). The values of *β*_1_ and *β*_2_ are equivalent in Eq. 1 and Eq. 3, and are constituted by a series of tree parameters which are all measurable and could be determined at a species-specific or any other relevant taxonomic level. In turn, *β*_3_ in theory can be approximated as 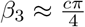 (where c is a unit conversion factor) under MST assumptions. In practise *β*_3_ does contain an additional term commonly referred to as a form factor, derived from stem tapering. As noted above we assume that area preservation occurs to generate our model for the height diameter relationship thus *β*_3_ should in theory have a value of 785.

The consequence of *β*_1*−*3_ being entirely set out by MST (see Eqs. 8 and 11) is that the allometric models are completely mechanistic, and thus can circumvent the need for statistical fitting procedures. The revised models retain many of the inferences of the original MST, given that their derivation is directly built upon the pre-existing theory. Note that the value *β*_2_ ∼ 0 can be asymptotically approached, which yields the original scaling relationships of 2/3 for *h* (Eq. 1) and 8/3 for *agb* (Eq. 3). Thus, Eqs. 1 and 3 are more generalized expressions of the MST scaling relationships, which become relevant only in tree species with traits that make *β*_2_ of relevant magnitude (e.g. small petioles and/or large proportion of vessels within them, see within methods). This explains the apparent lack of empirical fit observed for the original MST scaling in some cases (Muller-Landau et al., 2006; Picard et al., 2015; Swetnam et al., 2016; Jucker et al., 2017; Zhou et al., 2021). The resulting implication of the gMST modifier in Eqs. 1 and 3 compared with the original MST scaling is that it reduces the rate of increase in *h* and *agb* as the tree grows in *d*, thus being more relevant in larger trees.

## Model development and methods

### Theoretical basis for generalized Metabolic Scaling Theory (gMST) models and notation

Metabolic Scaling Theory (MST) models trees as a continually branching hierarchical network, running from the trunk to the petiole. The volume filling network produces a self-similar fractal (West et al., 1997, 1999) leading to relationships for the diameter (*d*_*k*_; m) and length (*l*_*k*_; m) of a parent branch (*k*) scaling from those of its daughter branch (*k* + 1):

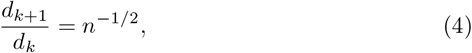

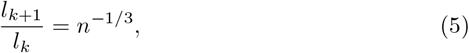

where *n* denotes the number of daughter branches that a parent branch splits into (typically *n* = 2), and *k* = 0, 1, …, *N* are the branching generations from the trunk (*k* = 0) to the petiole (*k* = *N*). MST postulates that area is preserved across all branching generations (Eq. 4), and that the system is an volume-filling network (Eq. 5). As the network is continuously branching, its total length 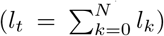 can be deduced through an infinitely scaling geometric series (West et al., 1999):

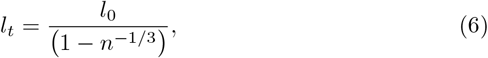

where *l*_0_ denotes the length of the trunk, and *l*_*t*_ is equivalent to the total height (*h*; m) of the tree. In our notation *d* is measured at breast height (*d*; m) is assumed to be equivalent to *d*_0_. Using these formulae, the *h* -*d* allometric equation can be derived as West et al. (1999):

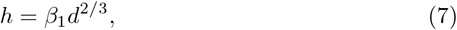

where *β*_1_ denotes a scalar:

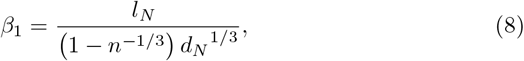

which becomes species-specific because it is regulated by tree parameters that can be determined at species-level: the length (*l*_*N*_), and diameter (*d*_*N*_) of the last branching generation (*N*), and the number *n* of daughter branches a species makes from each parent branch.

West et al. (1999) concluded that the *h* ∝ *d*^2*/*3^ scaling relationship derives from an area preserving and volume filling network as shown in Eqs. 4-5. They noted, along with previous literature (McMahon & Kronauer, 1976; Greenhill, 1881), that the 2/3 scaling exponent is an optimal mechanical relationship between *l*_*k*_ and *d*_*k*_ in order to resist buckling. By extension, the scaling exponent should be considered to apply to the area that gives mechanical structure. Consequentially, conductive area that does not generate mechanical strength should not necessarily scale to 2/3, and must be regulated by hydraulic limitations instead (K. Niklas, 2007). Moreover, key to MST is that an individual’s size scales with its resource distribution network (West et al., 1997). West et al. (1999) outlined how an optimum taper of conducting tissue within the branching network leads to a scaling relationship between the proportion of conductive tissue at each generation (*CA*_*k*_) and *d*_*k*_, such that *CA*_*k*_ ∝ *d*_*k*_^1*/*3^, with recent literature supporting the theory behind the predicted exponent (Koçillari et al., 2021; Rosell & Olson, 2019; Lechthaler et al., 2019). This tapering minimises energy expenditure for resource movement creating a hydraulic limitation to the potential maximum *h* that the tree could reach (hydraulic constraint). We postulate that this same limiting factor affects the mechanical limitations as well, because the proportion of conductive tissue within a cross-section of the trunk (*CA*_0_, which will hereby be referred to as *CA*) also increases as the tree grows in size, scaling to its *d* as *CA* ∝ *d* ^1*/*3^ (West et al., 1999). Since the mechanical limitation should only apply to the supportive tissue and the conductive tissue should play no part in supporting the branching network we propose that *h* ∝ *d* ^2*/*3^ scaling applies only to the proportion of tissue that has supportive function (*SA*, creating a mechanical constraint). This proposition effectively combines both the mechanical and hydraulic limitations into one, and it is the basis for the development of our generalized MST-based (gMST) allometry.

### Theory development for the generalized MST *h* -*d* allometric relationships

Alternatively to Eq. 7, we propose the true scaling relationship should give *h* as a function of only the proportion of *d* that supports the branching network. The proportion of supporting tissue, i.e. that primarily consisting of fibres, within a cross-section of the trunk (*SA*_0_, which will hereby be referred to as *SA*) should be entered within the scaling:

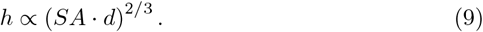

Moreover, MST also states that the proportion of conductive tissue *CA* within a given individual is proportional to *d*^1*/*3^, with this exponent derived from the rate of optimum vessel taper (West et al., 1999):

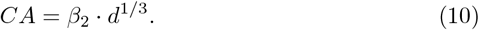

The variable *β*_2_ denotes a scalar that is determinable within MST (West et al., 1999, 1997) using the diameter of the final branching generation (*d*_*N*_), the number of conductive vessels within that petiole (*n*_*N*_) and the radii of each of these vessels (*a*_*N*_ ; m):

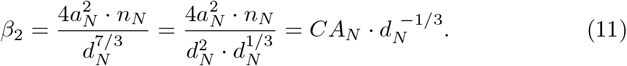

Alternatively *CA*_*N*_ which is the proportion of conductive material within a cross-section of the final branching generation (petiole) can be used to represent *β*_2_. Eq. 11 shows that *β*_2_ can be modelled as the ratio between *CA*_*N*_ and *d*_*N*_ ^1*/*3^, although the interpretation of the parameter is better considered with Eq. 10 showing that the ratios between the dimensions of trunk and petiole, *d* to *d*_*N*_, and their respective proportions of conductive (lumen) material, *CA* to *CA*_*N*_, scale together. Following Eq. 10, as *SA* = 1− *CA* (Fig. 1), the proportion of supportive material can be given as:

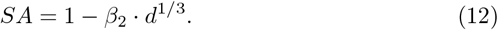

*SA* incorporates all types of tissue that can have supporting function within the trunk, comprising of any non-conductive tissue (non-lumen) (Ziemińska et al., 2013; Zanne et al., 2010). Applying Eq. 12 as a modifier to the scaling law therefore allows the interconnection between tissue types to be addressed within the model only using *d*, whereby only structurally relevant tissue is scaled to 2/3:

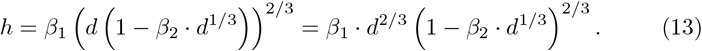

This gMST *h* -*d* allometric model (Eq. 1) based on a combination of MST scaling relationships includes the interconnection between the two tissue functional types: hydraulic constraints regulated by the tapering of conductive tissue and mechanical stability regulated by the proportion of supportive tissue, both bringing about a combined limitation to *h*. The mechanical constraint is predominant in large trees as they need an increasing *CA*, compared with smaller plants, to overcome hydraulic constrains to the upward flow of sap, and for that reason the *CA* scales itself to *d* as they grow (Eq. 10). Consequently, the effect of this component (Eq. 12) in the scaling relationship (Eq. 9) becomes more prominent for larger trees, which thus depart more from the original MST *h* -*d* 2/3 scaling law. On the other hand, Eq. 11 shows that the value of *β*_2_ depends on *a*_*N*_, *n*_*N*_ and *d*_*N*_, none of which can be physically null in a living individual and thus *β*_2_ must necessarily be a non-zero positive value (*β*_2_ ≠ 0). However, certain combinations of these tree parameters can overall make the proportion of conductive tissue in the petiole *CA*_*N*_ small, making *β*_2_ ∼ 0 negligible in practice (see Monte Carlo simulations in Methods), and thus the original MST *h* -*d* 2/3 scaling law approaches such cases well. Therefore, our allometric model in Eq. 13 is a generalized version of the original MST 2/3 scaling, rather than a substitution of it.

### Theory development for the generalized MST *agb* -*d* allo-metric relationships

Based on the same MST assumptions and constraints, West et al. (1999) also derived a 8/3 scaling law between the *d* of a tree and its biomass, and we can similarly propagate our gMST relationships to yield tree biomass. The aboveground biomass (*agb*; kg) of a cylindrical tree is calculated by the multiplication of its total aboveground volume (*v*_*tot*_; *m*^3^) and the average wood density (*ρ*; *gcm*^3^):

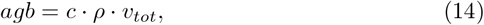

where *c* is a unit conversion factor (*c* = 10^3^). Eq. 14 works under the principle that *ρ* gives an average bulk density measurement, that takes into account both *CA* and *SA*. In theory one could assess the supportive volume and associated density, and thus estimate *agb*, although in practise taking such measurements is time consuming and labour intensive. It is simpler to consider the total volume, and average wood density (which is typically measured within our equations). Under the assumption of area preservation (Eq. 4), *v*_*tot*_ can be modelled as a cylinder (aggregated volume of the entire branching network *k* = 0 -*N* :

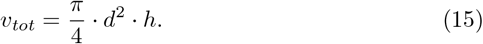

Yielding the subsequent *agb* equation,

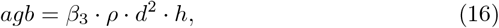

which is a *agb* ∼ *f* (*d, h*) allometric model (Eq. 2). *β*_3_ denotes a scalar which under the above-mentioned assumptions must approximate:

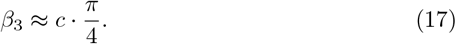

The unit conversion factor equals *c* = 10^3^ for the customarily used units stated above, which yields a value of 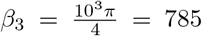. If centimetres are employed for *d* the overall factor becomes *c* = 10^*−*1^ which yields *β*_3_ = 0.0785. In comparison to real estimates this is a rather large value, with many authors assuming that the additional term (typically desribed as form factor, should be included) (Chave et al., 2014). It would therefore be unsurprising if this value is notably lower than 0.0785 when fitting.

While Eq. 16 provides a model to predict the *agb* of a tree from its measured *d* and *h* (*agb* ∼ *f* (*d, h*)) it is more typical to use a model dependent upon *d* only (*agb* ∼ *f* (*d*)), due to the difficulties of measuring height in the field. West et al. (1999) deducted the MST *agb* -*d* 8/3 scaling law by propagation of the *h* -*d* 2/3 scaling law. Similarly, the theoretical development of gMST can be propagated from *h* -*d* to yield the *agb* -*d* relationship by substituting Eq. 13 in Eq. 16, yielding:

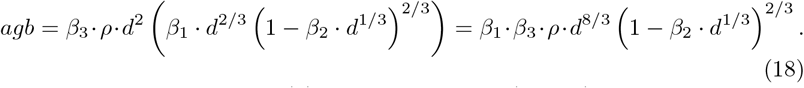

which is a gMST *agb* ∼ *f* (*d*) allometric model (Eq. 3). *β*_1_ and *β*_2_ can be directly propagated from the *h* -*d* equation, and thus retain their biophysical relevance (Eqs. 8 and 11). Thus, the inclusion of the component accounting for the scaling of supporting tissue (Eq. 9) also has a similar effect in modifying the original MST *agb* -*d* 8/3 scaling law between tree biomass and diameter, becoming more relevant for larger trees and those with traits resulting in large values of *β*_2_. The modifier concurrently allows for the original MST scaling law under *β*_2_ ∼ 0, and thus it is a generalized expression building upon the original MST 2/3 scaling, and not a substitution of it.

### Model fitting procedures

To test the validity of our proposed models, we contrasted them against a pantropical dataset compiled by Chave et al. (2014) and the Tallo allometry dataset (Jucker et al., 2022) estimating the values of *β*_1*−*3_ where appropriate. It has been argued that the maximum likelihood method (MLE) should be preferred for power-law estimations Stark et al. (2011). We decided to test both MLE and a Gauss-Newton algorithm for non-linear least squares (NLS) estimation, finding only marginal differences between them (Supplementary Table 1) and thus reporting MLE results only. Coefficients *β*_1*−*3_ were estimated using the ‘optim’ function for MLE and the ‘nls’ function for NLS, both within the ‘stats’ package in the R programming environment. In the case of MLE, the standard errors were estimated via bootstrapping 100 independent draws of 500 samples each.

The *h* -*d* model (Eq. 1) was fitted with initial values *β*_1_=100 and *β*_2_=0.2. No issues were found in terms of local minima, with the fitting process reaching the same result for wide ranges of initial values (any values of *β*_1_ ≥ 10 and *β*_2_ ≥ 0.01 up to *β*_2_ = 0.75 or larger depending on individual models). Our results were compared against those obtained by Chave et al. (2014)’s *h* -*d* model using their own compiled dataset, which in their case includes an environmental stress parameter (*E*) that explains a significant portion of variance in their model (with *d* specified in cm):

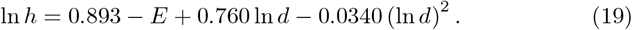

Thus, to give fairer comparison against Chave et al. (2014), we made a version of the gMST model that incorporates *E* in a similar manner (i.e. converted to original scale):

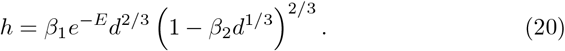

Furthermore, the Tallo dataset (Jucker et al., 2022) contains data on nearly half a million individual trees with measured *d* and *h* sufficient to derive separate *h* -*d* models according to functional type - gymnosperms and angiosperms - and ecoregion. In some cases, fitting these models required more refined initial guesses. Environmental stress parameters were found for each tree using the computeE function in the BIOMASS package (Réjou-Méchain et al., 2017), although less than 25% had a valid output, thus environmental stress was not used in association with the Tallo dataset. An F-test for nested models was used to evaluate the significance of all these stratum-wise gMST models against the simpler 2/3 scaling (Table 2).

The *agb* models (Eqs. 2-3) were tested only against Chave et al. (2014) data, because it incorporated all the predictors needed (*ρ, d* and *h*). In this case we tested two groups of models: those predicting *agb* from both *d* and *h* (*agb* ∼ *f* (*d, h*)), and those predicting from *d* only (*agb* ∼ *f* (*d*)). All the *agb* models were developed by directly propagating the 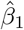 and 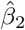 values from the fitted *h* -*d* results, and *β*_3_ was estimated via MLE. For comparison, these were tested against Chave et al. (2014) *agb* ∼ *f* (*d, h*) model using their own data (with *d* specified in cm):

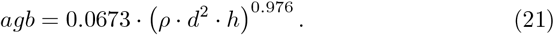

Our results were also compared against Chave et al. (2014)’s *agb* ∼ *f* (*d*) model, which involves the use of *E* (with *d* specified in cm):

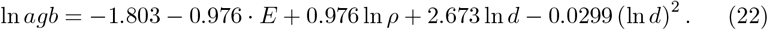

And again, we made a version of the gMST model that incorporates E for fair comparison:

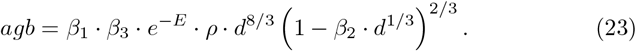

The initial value for the MLE estimation was *β*_3_ = 785, again with a wide range of initial values of around *β*_3_ = 50 − 3, 000 reaching identical solutions.

### Monte Carlo simulations

To deduce whether the estimated 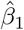 and 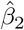 values are realistic in natural systems, and reflect on the biological relevance of these coefficients, Eqs. 8 and 11 were used within Monte Carlo simulations to obtain frequency distributions for *β*_1_ and *β*_2_ that derive from plausible values of the tree parameters (*d*_*N*_, *n*_*N*_, *a*_*N*_, *l*_*N*_ and *n*). These parameters are tree architecture traits (e.g. dimensions of the stem and petiole). Some of the simulated values were deducted from the literature, whereas others followed logical inter-dependencies among these traits (such as *d*_*N*_ and *l*_*N*_). We assumed a log-uniform distribution within a range of *l*_*N*_ values from 3 mm to 1 m (Poorter & Rozendaal, 2008). The number of daughter branches that a parent branch splits into was assumed to follow a geometric distribution truncated to a range *n* = 2 − 6. Since some of the tree traits are mutually inter-dependent, as *d*_*N*_ invariably changes with *l*_*N*_ and *n*, it would be unrealistic to draw random independent values from each of them. For this reason, we modelled their relationship through the rearrangement and combination of Eqs. 4-7:

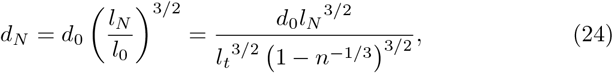

which follows from the relationship between the ratios of *d*_*N*_ to *d*_0_ and *l*_*N*_ to *l*_0_, i.e. the dimensions of the first and last branching generations, as determined by MST assumptions (Eqs. 4-5). Then, by solving *l*_0_ in Eq. 6, it was substituted by *l*_*t*_ and *n* in the denominator, meaning a function could be reached for *d*_*N*_ ; *d*_*N*_ ∼ *f* (*l*_*N*_, *n, d*_0_, *l*_*t*_) in Eq. 28, where *l*_*N*_ and *n* have been simulated in previous steps and paired values of *d*_0_ and *l*_*t*_ can be retrieved from actual empirical samples (n.b. *d*_0_ ≡ *d* and *l*_*t*_ = *h*). In a Monte Carlo simulation with a sample of size 1,000, we drew independent samples from the pantropical (Chave et al., 2014) and Tallo datasets (Jucker et al., 2022), plus from the above-mentioned assumed distributions for *l*_*N*_ and *n*, to generate a distribution of *d*_*N*_ values associated with them. This provided a set of realistic paired values for *d*_*N*_, *l*_*N*_, and *n* which were combined through Eq. 8 to derive plausible *β*_1_ values under natural conditions.

Following up from the paired values obtained for *β*_1_, a similar procedure was employed to derive a distribution of plausible joint *β*_2_ values under natural conditions. *β*_2_ can be shown as a function of *d*_*N*_ and *CT*_*N*_ (*β*_2_ ∼ *f* (*d*_*N*_, *CT*_*N*_)). Again, the relationship of these dimensions in the petiole and the trunk, *d*_*N*_ to *d*_0_ and *CT*_*N*_ to *CT*_0_, can be related via MST assumptions (Eq. 11):

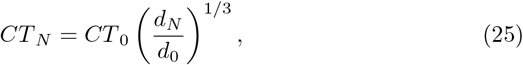

unfortunately, datasets with *CT*_0_ measurements along with their respective *d*_0_ measurements are not available. To overcome this difficulty, we simulated *CT*_0_ values assuming a non-central Beta distribution *CT*_0_ *NonCentralBeta* (*α* = 2, *β* = 20, *δ* = 0.1), which we regarded to faithfully model the lumen proportions observed in the literature (Ziemińska et al., 2013; Zanne et al., 2010). The resulting plausible distributions for *β*_1_ and *β*_2_ are shown in Figs. 3A-B, where the 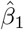 and 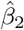 values estimated for empirical models are shown (table 1 in results) to illustrate that those values correspond well to the combination of realistic assumptions in the tree parameters considered in Eqs. 8 and 11. An ellipse of 95% confidence intervals (CIs) for *β*_1_-*β*_2_ combinations was derived from a *χ*^2^ distribution with 2 degrees of freedom (Figs. 3A-B). Since any *β*_1_-*β*_2_ combination within that ellipse provides 95% CIs for the model, we chose to determine those that would correspond to the tallest and shortest tree with quadratic mean diameter in the dataset. In order to achieve that, we calculated the height that would correspond to quadratic mean diameter, for every *β*_1_-*β*_2_ combination calculated in that ellipse, and from those results we extracted values corresponding to the minimum and maximum tree height, which are those shown for the gMST *h* -*d* models in Figs. 3C-D. These were also propagated to the gMST *agb* -*d* model in Fig. 4C-D.

**Table 1:** Results of height-diameter modelling across the Chave et al. (2014) (abbreviated to Ch) and the Tallo (abbreviated to Tll Jucker et al. (2022)) datasets, with the original 2/3 MST scaling given across the entirety of both datasets for comparison. Chave *et al*. (2014)’s height diameter model is also given for comparison against the gMST formulations with and with environmental stress (*E*) included. The Tll dataset has been divided to give overall angiosperm and gymnosperm models, and further stratified to give angiosperm and gymnosperm models for each ecoregion that the dataset allows.

|  | Coef ests |  |  |  | Model eval |  |  |  |  |  |
| --- | --- | --- | --- | --- | --- | --- | --- | --- | --- | --- |
| | $\hat{\beta}_1$ | $\hat{\beta}_1\text{SE}$ | $\hat{\beta}_2$ | $\hat{\beta}_2\text{SE}$ | $R^2$ | RMSE | RMSE (%) | MD | MD (%) | n |
| Ch |  |  |  |  |  |  |  |  |  |  |
| 2/3 scaling | 44.17 | 0.17 | - | - | 0.73 | 5.59 | 34.87 | -0.31 | -1.92 | 4,004 |
| Ch h-d model | - | - | - | - | 0.87 | 3.82 | 23.83 | -0.39 | -2.45 | 4,004 |
| gMST with E | 66.73 | 0.44 | 0.53 | 0.01 | 0.88 | 3.80 | 23.71 | -0.07 | -0.41 | 4,004 |
| gMST without E | 52.04 | 0.53 | 0.28 | 0.02 | 0.74 | 5.52 | 34.44 | 0.11 | 0.67 | 4,004 |
| TII |  |  |  |  |  |  |  |  |  |  |
| 2/3 scaling (whole TII dataset) | 37.25 | 0.25 | - | - | 0.51 | 6.14 | 49.73 | -0.36 | -2.96 | 496,822 |
| gMST (whole TII dataset) | 45.10 | 0.94 | 0.34 | 0.09 | 0.52 | 6.07 | 49.20 | -0.02 | -0.16 | 496,822 |
| Angio 2/3 scaling | 36.21 | 0.31 | - | - | 0.49 | 6.13 | 53.11 | -0.48 | -4.19 | 333,285 |
| Angio gMST | 44.77 | 0.92 | 0.37 | 0.05 | 0.51 | 6.04 | 52.35 | -0.08 | -0.66 | 333,285 |
| Gymno 2/3 scaling | 34.72 | 0.20 | - | - | 0.50 | 5.52 | 44.68 | 0.11 | 0.86 | 102,670 |
| Gymno gMST | 30.68 | 1.17 | -0.30 | 0.67 | 0.50 | 5.51 | 44.60 | 0.00 | 0.01 | 102,670 |

**Figure 2:**
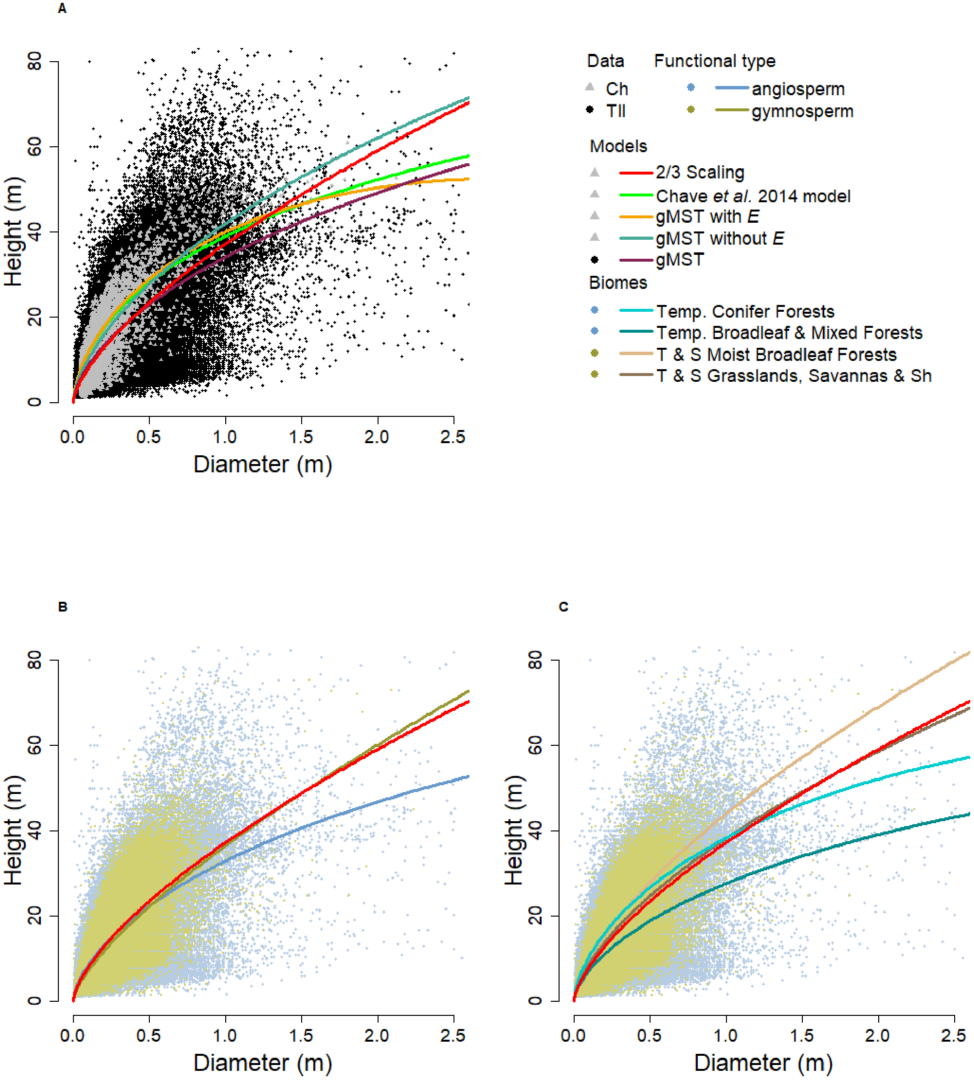
The figure gives the plotting of a number of the height-diameter models. A) shows the comparison of Chave et al. (2014) against gMST (orange line includes E, blue line does not include E) models, the original MST 2/3 scaling for the Chave et al. (2014) dataset in red and Chave et al. (2014)’s model is given in green. Both datasets are given, black represents the Tallo dataset (abbreviated Tll) (Jucker et al., 2022), and the Chave et al. (2014) data is shown in grey. B) only shows the Tll data, and shows the scaling of angiosperms and gymnosperms, as well as the original 2/3 MST scaling and C) shows the Tll dataset against a selection of four ecoregion models: Temperate conifer Forests - gymnosperm, Temperate Broadleaf & Mixed Forests - gymnosperm, Tropical & Subtropical Moist Broadleaf Forests - angiosperm, Tropical & Subtropical Grasslands, savannas & Shrublands - angiosperm, compared against the 2/3 scaling for the Tll dataset. A small fraction of the Tll dataset is not given on the plots, with the largest individual tree being about 8m wide, thus the axis limits are given to improve graph readability. Within the legend Tropical and subtropical has been abbreviated to T & S, and Shrublands has been abbreviated to Sh.

**Figure 3:**
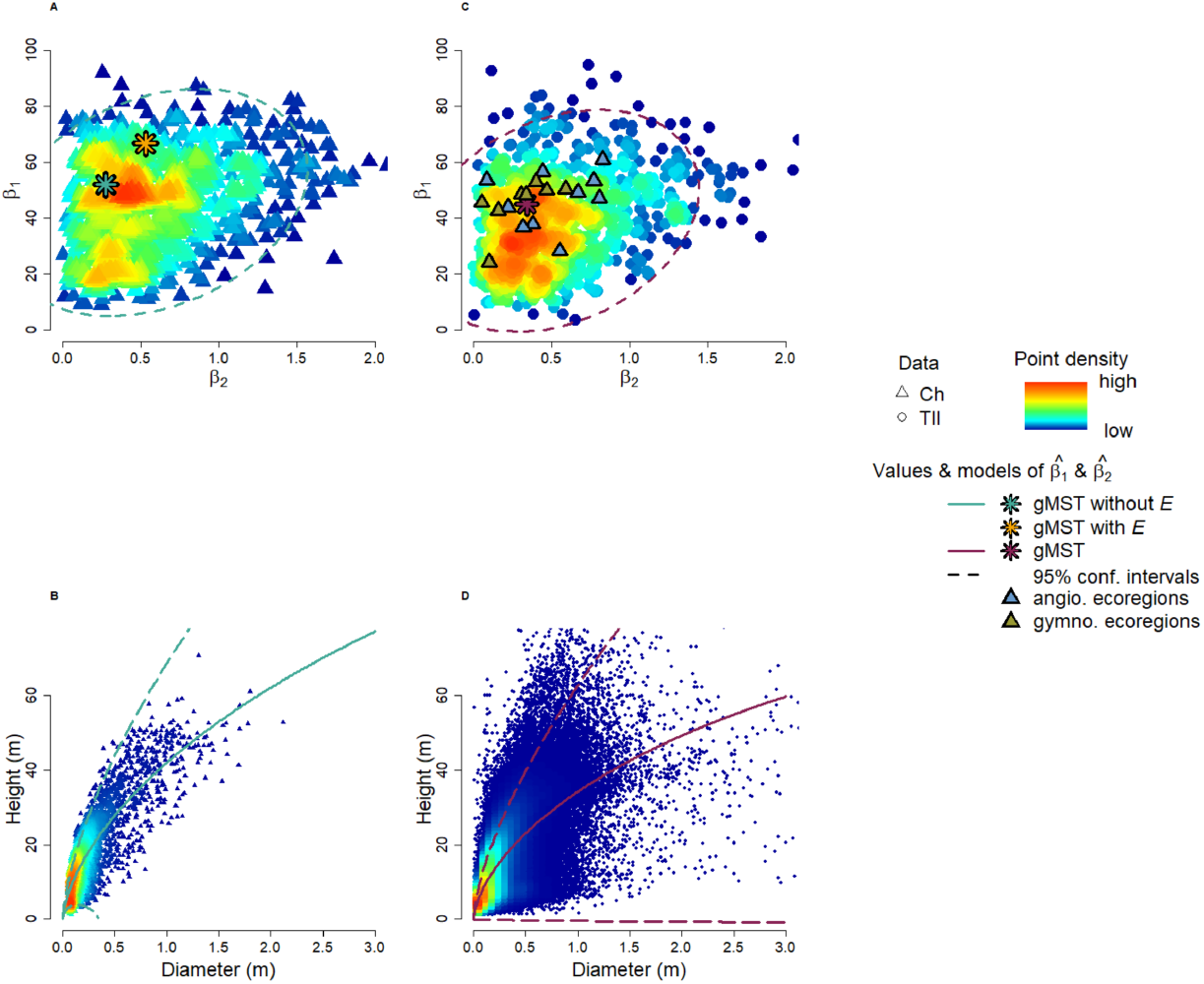
The figure gives the results of the Monte Carlo simulations for the height-diameter modelling, showing the values of *β*_1_ and *β*_2_ generated from combinations of plausible trait (*d*_*N*_, *n*_*N*_, *a*_*N*_, *l*_*N*_ and *n*) values (A and C). The values of 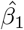 and 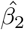generated from model fitting across are also given for reference. A) gives the plausible *β*_1_ and *β*_2_ values for Chave et al. (2014) with the predicted parameter values from the gMST fitting being shown (orange represents the model including E, and green represents the model without E). C) gives the plausible values for the Tll dataset (Jucker et al., 2022), with the overall fitted *β*_1_ and *β*_2_ values given as well as the ecoregion specific values being plotted. 95% confidence intervals are also given for reference. B) and D) show how the 95% confidence intervals affect the overall scaling for each dataset (B for Chave et al. (2014) and D for Tll (Jucker et al., 2022)).

**Figure 4:**
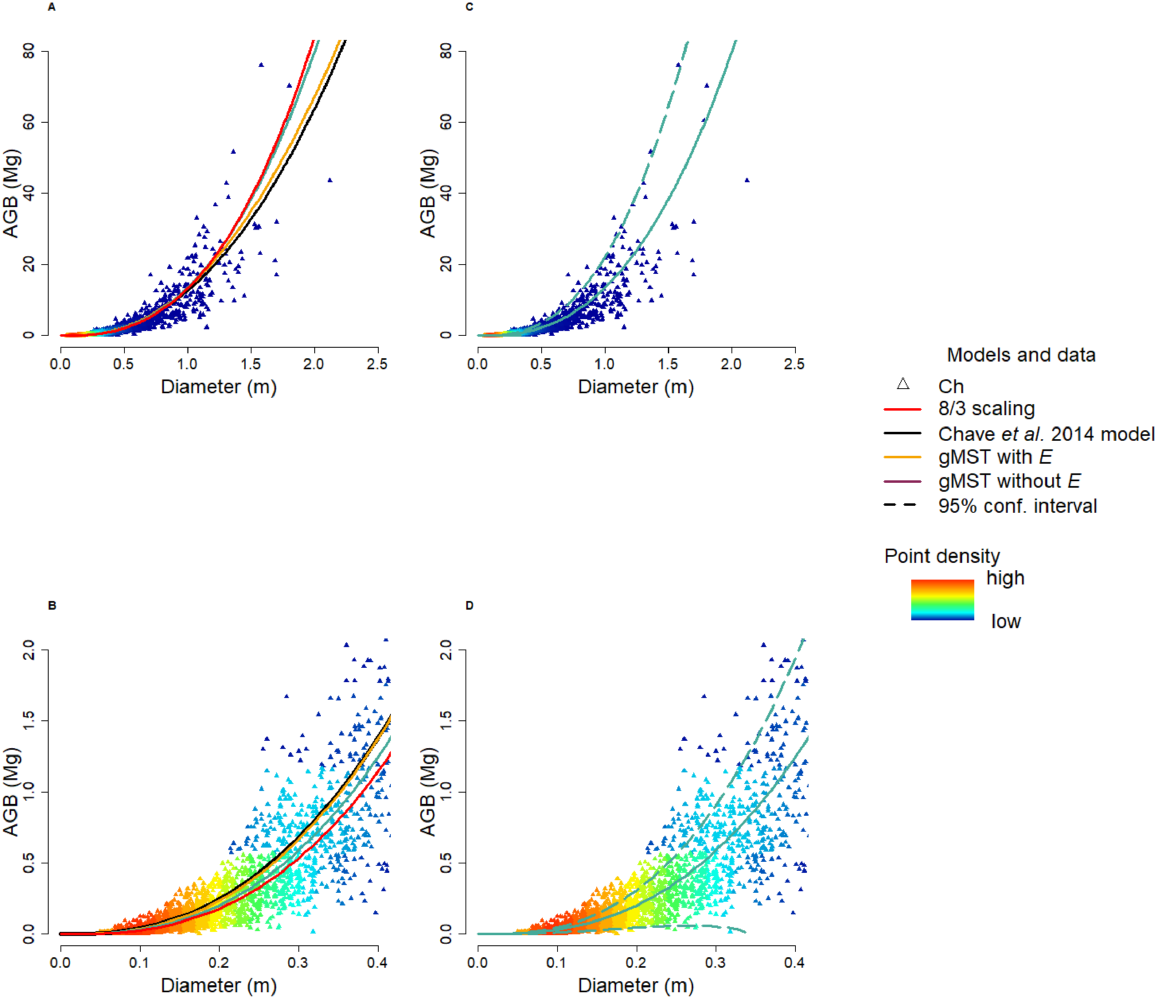
A) shows the gMST fitted models with and without *E*, against the orginial 8/3 *agb* MST scaling. C) gives the 95% confidence intervals produced through the propagation of the *β*_1_ and *β*_2_ 95% values. B) and D) then give finer scale figures to improve readability at low values of diameter and *agb*. Chave et al. (2014) has been abbreviated to Ch.

## Results

Since our generalized MST models are mechanistically derived, in principle predictions for individual trees could be generated from tree parameters (i.e. *d*_*N*_ to *d*_0_ and *l*_*N*_ to *l*_0_) using Eqs. 8, 11 and 20. However, there is no such type of data available to prove the global validity of our models in respect to Eqs. 8, 11 and 20, but statistically-fitted values of 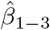 can be obtained through global allometry datasets (see model fitting procedures in Methods). The models can thus be compared against widely used statistical models (e.g Chave et al. (2014)), and the mechanistic significance of the resulting *β*_1_-*β*_2_ values can be evaluated in comparison to the Monte Carlo simulations (Eqs. 8, 11 and 20, see Monte Carlo simulations in Methods). To test our gMST models, we carried out statistical fits against worldwide data compiled for pantropical (Chave et al., 2014) and global allometry testing (Jucker et al., 2022). Estimated values for coefficients 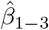 in Eqs. 1-3 were fitted using these datasets, with stratifications based on tree functional type and ecoregion across the global Tallo dataset (sensu Jucker et al. (2022)). Furthermore, to give a fair comparison between the gMST models and the models created by Chave et al. (2014), the environmental stress parameter (*E*) generated by Chave et al. (2014) was integrated into a subset of our gMST models (see model fitting procedures in Methods). The *E* parameter combines factors that explain hydraulic constrains to tree growth, namely the temperature and precipitation seasonality and climatic water deficit.

### Height-diameter model

Table 1 and 2 show the results of the *h* -*d* models fit through maximum likelihood estimation (MLE) (see model fitting procedures in Methods), including estimates and standard errors for 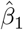 and 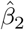, as well as their respective coefficients of determination (*R*^2^) root mean squared error (RMSE) and mean differences (MD). The 2/3 scaling relationship originally predicted by MST showed a clear lack of fit to the data, with a bias most prevalent for larger trees, especially for angiosperms (Table 1 plus S.table 1, and Fig. 2). For this reason, the prevalent pantropical allometric model derived by Chave et al. (2014) diverges significantly from the 2/3 scaling rule.

**Table 2:** Biome specific results for our proposed gMST model, giving coefficient values, model evaluation metrics and the results of a series of ANOVA tests, performed between the original 2/3 scaling and our gMST model. Tropical and subtropical has been abbreviated to T & S, and Shrublands has been abbreviated to Sh within the table. The ANOVA tests were carried out upon the nls models, that proved almost identical to the maximum likelihood estimation results (see supplementary table). The maximum likelihood estimation coefficients are however given for each biome in the table presented. Difficultly in the nls fitting upon the Deserts and Xeric Shrubland lead to NA values for the ANOVA tests.

|  | Coef ests |  |  |  | Model eval |  |  |  |  | ANOVA results |  |  |
| --- | --- | --- | --- | --- | --- | --- | --- | --- | --- | --- | --- | --- |
| | $\hat{\beta}_1$ | $\hat{\beta}_1$SE | $\hat{\beta}_2$ | $\hat{\beta}_2$SE | R<sup>2</sup> | RMSE | RMSE (%) | MD | MD (%) | n | F value | p value |
| Angio |  |  |  |  |  |  |  |  |  |  |  |  |
| T & S Moist Broadleaf Forests | 56.59 | 0.72 | 0.44 | 0.04 | 0.72 | 4.62 | 35.27 | 0.17 | 1.31 | 128305 | 9,666.83 | 0 |
| T & S Dry Broadleaf Forests | 47.24 | 0.75 | 0.80 | 0.09 | 0.52 | 2.15 | 34.05 | -0.12 | -1.92 | 27813 | 5,619.93 | 0 |
| Temp. Broadleaf & Mixed Forests | 44.07 | 1.23 | 0.22 | 0.25 | 0.60 | 6.21 | 48.18 | 0.18 | 1.37 | 96222 | 1,337.23 | 0 |
| Temp. Conifer Forests | 39.17 | 1.00 | -0.13 | 0.13 | 0.62 | 5.75 | 48.72 | -0.16 | -1.33 | 9228 | 9.81 | 0 |
| Boreal Forests/Taiga | 53.74 | 1.18 | 0.09 | 0.13 | 0.59 | 3.90 | 28.27 | 0.02 | 0.18 | 1441 | 0.3 | 0.58 |
| T & S Grasslands, Savannas & Sh | 38.02 | 0.77 | 0.38 | 0.09 | 0.55 | 4.12 | 46.25 | -0.01 | -0.06 | 15749 | 338.42 | 0 |
| Temp. Grasslands, Savannas & Sh | 49.27 | 0.73 | 0.67 | 0.04 | 0.63 | 3.83 | 43.30 | 0.13 | 1.50 | 5588 | 613.94 | 0 |
| Flooded Grasslands & Savannas | 23.36 | 0.61 | -0.64 | 0.36 | 0.69 | 2.65 | 42.50 | 0.05 | 0.72 | 443 | 4.35 | 0.04 |
| Montane Grasslands & Shrublands | 36.79 | 0.46 | 0.32 | 0.03 | 0.62 | 4.53 | 37.22 | -0.03 | -0.21 | 3701 | 84.73 | 0 |
| Tundra | 53.49 | 0.69 | 0.77 | 0.03 | 0.74 | 2.38 | 29.48 | 0.08 | 1.01 | 336 | 26.98 | 0 |
| Mediterranean Forests, Woodlands & Sh | 28.31 | 0.85 | 0.55 | 0.05 | 0.10 | 5.53 | 64.80 | -0.36 | -4.28 | 43323 | 1,985.32 | 0 |
| Deserts & Xeric Shrublands | 3.06 | 0.31 | -31.60 | 2.79 | 0.46 | 3.43 | 76.22 | -0.01 | -0.22 | 380 | NA | NA |
| Mangroves | 61.02 | 0.44 | 0.83 | 0.01 | 0.70 | 3.51 | 32.21 | 0.18 | 1.69 | 739 | 321.7 | 0 |
| Gymno |  |  |  |  |  |  |  |  |  |  |  |  |
| T & S Moist Broadleaf Forests | 50.04 | 0.43 | 0.47 | 0.01 | 0.77 | 3.96 | 29.07 | 0.10 | 0.75 | 653 | 59.81 | 0 |
| T & S Coniferous Forests | 13.25 | 0.13 | -3.83 | 0.09 | 0.90 | 2.44 | 13.49 | 0.03 | 0.18 | 105 | 27.28 | 0 |
| Temp. Broadleaf & Mixed Forests | 42.87 | 1.02 | 0.16 | 0.24 | 0.56 | 5.25 | 36.28 | 0.09 | 0.65 | 24128 | 40.07 | 0 |
| Temp. Conifer Forests | 45.63 | 1.17 | 0.05 | 0.17 | 0.77 | 4.99 | 33.41 | 0.37 | 2.48 | 17024 | 15.42 | 0 |
| Boreal Forests/Taiga | 48.58 | 0.96 | 0.31 | 0.08 | 0.68 | 3.24 | 27.02 | 0.11 | 0.96 | 9640 | 77.39 | 0 |
| T & S Grasslands, Savannas & Sh | 50.44 | 0.58 | 0.59 | 0.02 | 0.64 | 2.33 | 21.05 | 0.01 | 0.11 | 972 | 64.54 | 0 |
| Temp. Grasslands, Savannas & Sh | 48.36 | 0.79 | 0.34 | 0.05 | 0.68 | 3.37 | 31.79 | 0.03 | 0.26 | 3624 | 50.33 | 0 |
| Montane Grasslands & Shrublands | 53.17 | 0.74 | 0.40 | 0.03 | 0.59 | 6.13 | 33.14 | 0.16 | 0.87 | 8336 | 224.41 | 0 |
| Tundra | 30.34 | 0.72 | -0.58 | 0.21 | 0.71 | 2.51 | 36.94 | 0.00 | 0.02 | 484 | 3.83 | 0.05 |
| Mediterranean Forests, Woodlands & Sh | 24.33 | 0.62 | 0.10 | 0.42 | 0.49 | 2.72 | 31.06 | 0.02 | 0.22 | 37375 | 21.03 | 0 |
| Deserts & Xeric Shrublands | 16.66 | 1.06 | -2.82 | 0.86 | 0.68 | 4.98 | 31.48 | 0.11 | 0.71 | 328 | 11.81 | 0 |

Our *h* -*d* gMST model outperformed both the 2/3 scaling rule and the pantropical model of Chave et al. (2014) (Table 1). These gave slightly biased mean predictions of tree heights (MD = -0.31 m and -0.39 m), which were improved to just -0.07 m by the gMST model. Additionally, the RMSE of 3.80 m obtained by gMST was also an improvement compared to the 2/3 scaling and even to the 3.82 m RMSE obtained by the model fitted by Chave et al. (2014) with that same data. This implies that the model form proposed in Eq. 1 is most appropriate, and betters both the simpler 2/3 scaling relationship and the quadratic model form (on a logarithmic scale) chosen by Chave et al. (2014).

The deviation from the 2/3 scaling law was more notable in angiosperms than in gymnosperms, both overall and within specific ecoregions (Fig. 2B and Fig. 2C). This effect could be recognized through the marked difference in the estimates for 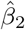 (Table 1 and 2). Gymnosperms models tended to exhibit lower 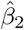 values than angiosperms. When considering functional type and ecoregion combinations (Fig. 2C, table 1, table 2 and S.Fig. 1 and S.Fig. 2 in the supplementary material), the majority of the parameters proved to be significant (*p <* 0.05, calculated through NLS) apart from three 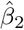 values for gymnosperm ecoregion models and two 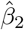 angiosperm ecoregion models (table 2). All but two models proved significantly different from the original 2/3 scaling being the Boreal Forest/Taiga ecoregion for angiosperms and the Tundra ecoregion for gymnosperms (table 2). Interestingly a number of the 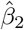 estimates were negative (table 1, table 2 and S.table 1 in the supplementary material), which violates the theory set out above. Only one of these values were significant across the ecoregions, but the three insignificant negative gymnosperm values are likely to have influenced the overall 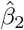 value for gymnosperm data. The negative value for all gymnosperm species is therefore a bad representation of the whole dataset, although is significant.

Both *β*_1_ and *β*_2_ can be considered as a function of tree parameters (i.e. *d*_*N*_ to *d*_0_ and *l*_*N*_ to *l*_0_), given Eqs. 8, and 11. To investigate the plausibility of the statistically fitted values obtained in Table 1 and 2, we carried out Monte Carlo simulations using realistic values of the tree parameters involved at each of them, to generate biologically plausible joint distributions of *β*_1_ and *β*_2_. The results (Figs. 3A-B) demonstrate that the values 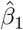 and 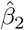 obtained empirically (Table 1 and 2) are biologically plausible, as they conform with realistic values of tree parameters in Eqs. 8 and 11. Moreover, Figs. 3C-D give the model trajectories generated from 95% confidence intervals of the Monte Carlo simulations, showing that the majority of the field observations are contained within these intervals.

### Biomass-Diameter model

The *agb* -*d* models (Eqs. 2-3) were tested against Chave et al. (2014)’s dataset (abbreviated to Ch within the tables), because it was the only dataset that incorporated all the data needed (*ρ, d, h* and importantly *agb* data). Prior to reflecting on the *agb* ∼ *f* (*d*) relationship, it is noteworthy to consider the relationships using both tree height and diameter as predictors for aboveground biomass (*agb* ∼ *f* (*d, h*)). Chave et al. (2014) widely used pantropical model equivalent gMST derived equation, with the form *agb* = *β*_3_ (*ρd*^2^*h)* (Eq. 2). The *agb* = 0.0673 ·(*ρd*^2^*h*)^0.976^ (n.b. diameter given in cm) was compared against the theoretical gMST model achieved a better RMSE than the statistical fit of Chave et al. (2014), although proved to be more biased. The *agb* ∼ *f* (*d, h*) models are limited by the requirement of *h* measurements, which are often unavailable (Valbuena et al., 2016; Feldpausch et al., 2012) and thus the validity of the gMST *agb* ∼ *f* (*d*) relationship was considered further.

The gMST allometry for *agb* ∼ *f* (*d*) (Eq. 3) was employed by propagating values of 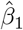 and 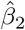 from each associated *h* ∼ *f* (*d*) model (Table 1), and subsequently fitting *β*_3_. Interestingly the gMST model excluding *E* performed best out of our new models, proving to have a lower RMSE than the widely used pantropical models (Chave et al., 2014), although the bias proved to be higher.

Improvements upon the *R*^2^ was also observed. 95% confidence intervals for this model (Figs. 4B and 4D) were generated by propagating the Monte Carlo simulation results for *β*_1_ and *β*_2_ (Figs. 3A-B) to the *agb* ∼ *f* (*d*) equation, again demonstrating that the realistic variation in *β*_1_ and *β*_2_ adequately encompasses the majority of field observations. The best performing *agb* ∼ *f* (*d*) model (in terms of *R*^2^ and RMSE) proved to be that of the original 8/3 scaling, although it was more biased (MD = -0.030) than Chave et al. (2014) and the gMST without E model. The comparison of the *agb* ∼ *f* (*d*) models is shown in Fig. 4A illustrates that the model trajectories have only slight differences. Thus, generally speaking the choice of *agb* -*d* model matters less than for *h* -*d*, although one must consider the uncertainty in terms of relative error implied by the use of these models (RMSE = 10.8-49.8 % for *h*, and RMSE = 102.8-123.8 % for *agb*).

## Discussion

Our results demonstrate that the gMST relationships are unbiased and show a significant level of empirical fit to global tree datasets that make them universally applicable. We also show that the statistical fit to data yields coefficient values that are plausible for realistic magnitudes of the tree parameters involved in Eqs. 8 and 11, demonstrate the potential biological relevance of our gMST. The main asset of the gMST models is their mechanistic derivation underpinned by MST assumptions. Remarkably, the theory based gMST models outperformed the statistical models of Chave et al. (2014), in terms of *h* -*d* modelling, using their complied dataset and proved comparable when modelling *agb*. These results support MST itself, resolving the criticism pertaining to the lack of empirical support to MST scaling for large trees and tropical areas (Muller-Landau et al., 2006; Picard et al., 2015; Swetnam et al., 2016; Jucker et al., 2017; Zhou et al., 2021). Furthermore, gMST concurrently allows for the original MST scaling law (*β*_2_ ∼ 0), explaining why the original MST scaling relationships fit adequately in some scenarios.

The gMST *h* -*d* model proposed (Eq. 1) performed exceptionally well, demonstrating a marked improvement upon both the original 2/3 scaling law and the widely used pantropical model by Chave et al. (2014) (Table 1 and Fig.2). It was superior to the 2/3 scaling against the Tallo dataset(Jucker et al., 2022), especially for angiosperms, and to a slightly lesser extend gymnosperms (Tables 1 and Tables 2). Moreover, the simple *agb* ∼ *f* (*d, h*) model based on MST assumptions (Eq. 2) outperformed Chave et al. (2014)’s pantropical model (Table 3). The propagation of the gMST *h* ∼ *f* (*d*) model to derive Eq. 3 (*agb* ∼*f* (*d*)) gave results comparable to the original 8/3 scaling law and Chave et al. (2014)’s model (Table 3 and Fig. 4). Nonetheless it should be noted that at the *agb* -*d* level our study could only test against the pantropical dataset. Global data including *agb* measured through destructive sampling is needed to clarify functional type and ecoregion dependencies for the *agb* -*d* relationship. We also recommend that further research focuses on acquiring empirical measurements of the tree parameters involved in Eqs. 8 and 11, because once sufficient data becomes available the gMST models could potentially become grounds for determining tree biomass in the field and circumvent the need for destructive sampling.

**Table 3:** Results for *agb* related models, giving the comparison of equations utilising both height and diameter as well as models purely using diameter. The fitted values of 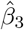 are given within the table for the associated models. E denotes environmental stress as calculated by Chave et al. (2014)

|  | Coef ests |  | Model eval. |  |  |  |  |
| --- | --- | --- | --- | --- | --- | --- | --- |
| | $\hat{\beta}_3$ | $\hat{\beta}_3$SE | R<sup>2</sup> | RMSE | RMSE (%) | MD | MD (%) |
| Ch f(d,h) | - | - | 0.91 | 1.21 | 106.39 | 0.00 | -0.20 |
| f(d,h) | 527.00 | 2.18 | 0.91 | 1.17 | 102.78 | 0.04 | 3.59 |
| Ch f(d) | - | - | 0.87 | 1.40 | 123.83 | 0.00 | -0.27 |
| 8/3 f(d) | 472.76 | 1.96 | 0.89 | 1.30 | 114.64 | -0.03 | -2.62 |
| gMST f(d) with E | 526.50 | 2.82 | 0.88 | 1.35 | 119.00 | 0.03 | 2.80 |
| gMST f(d) without E | 507.65 | 2.28 | 0.89 | 1.30 | 114.93 | 0.01 | 0.67 |

A clear difference between tree functional types was evident within the 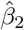 results (Table 1, Table 2 and Figs. 2B-C), likely due to divergences at the cellular level. Disparities in the tree parameters influencing *β*_2_ (Eq. 11) have indeed been observed between angiosperms and gymnosperms (Sperry et al., 2008, 2006). Angiosperms have differentiated cell types, vessels and fibres, each dedicated to conductive and supportive functions respectively. This tissular differentiation brings about a functional separation at different areas within the cross-section of the tree stem, which other authors have referred to as lumen and non-lumen fractions (for *CA* and *SA* respectively)(Ziemińska et al., 2013; Zanne et al., 2010). Alternatively, gymnosperms have integrated conductive and supportive functionality in their tissues through tracheids, and this functional integration of tissue leads to greater deviations from Murray’s law for gymnosperms than angiosperms (McCulloh et al., 2004). Thus, the limitation in the *h* of a tree is more regulated by mechanical constraints in gymnosperms (MST), while angiosperms tend towards hydraulic constraints (Eq. 10 in Methods), with gMST effectively integrating both controls, explaining why a number of the gymnosperm *β*_2_ proved insignificant and negative.

There were notable levels of variation in the estimated 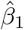 and 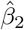 values across ecoregions, which gives clues on their biological relevance. All values were included within the distributions of plausible coefficient values, which we derived from reasonable values of tree parameters (Eqs. 8 and 11) via Monte Carlo simulations (Fig. 4). Environmental factors are likely to explain the variance across ecoregions in species of the same functional type, with *β*_1_ and *β*_2_ changing in accordance to plant survival strategies, given that tree parameters (such as vessel dimensions) change according to the environment (Preston et al., 2006). Stratum-wise estimations of 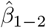 therefore reflect the combinations of optimal tree parameters for the growth and survival of a given species in its typical habitat. This idea is also reinforced by the high proportion of explained variability added by *E* in the gMST *h* -*d* model. It should be pointed out that our use of *E* within the gMST models (Eqs. 20 and 23 in Methods) is contentious, given the calculation of *E* is intrinsically linked to models created by Chave et al. (2014). Further research should be devoted to find the mechanistic integration of the factors intrinsically included in *E* – temperature and precipitation seasonality, and water deficit – in the gMST model: how they affect the tree parameters involved in *β*_1_ and *β*_2_ (Eqs. 8 and 11) Fortin et al. (2019), given that specific species traits could compensate for the hydraulic constraints on tree growth that these factors impose.

## Conclusions and future perspectives

We present new generalised equations through the consideration of differentiated tissue functions. These equations are characterised by theory-determinable exponents and coefficients, allowing for a fully generalised and globally applicable model form. The exponents are fixed by MST scaling, whereas the coefficients are defined by tree traits that vary with ecoregion and tree functional type, likely driven as adaptations to different climatic conditions. We showed that our equations outperform statistically fitted models that are widely used for large scale *h* estimations. Our gMST models should provide a new avenue of scientific development towards global theory-based models for tree allometry, giving a possibility to mechanistically link histological and macroscopic traits. They will also pave the way for large scale, accurate *agb* and forest carbon estimation globally, utilising new methods and technology such as satellite lidar available through the Global Ecosystem Dynamics Investigation (GEDI) mission.

## Acknowledgements

The authors would like to thank the National Environmental Research Council (NERC) for funding Stuart Sopp’s PhD, and declare that there are no conflicting interests.

